# Emerging Tumor Development by Simulating Single-cell Events

**DOI:** 10.1101/2020.08.24.264150

**Authors:** Jakob Rosenbauer, Marco Berghoff, Alexander Schug

**Affiliations:** Jülich Supercomputing Centre, Forschungszentrum Jülich, Wilhelm-Johnen-Straße, 52428 Jülich, Germany; Steinbuch Centre for Computing, Karlsruhe Institute of Technology, Kaiserstraße 12, 76131 Karlsruhe, Germany; Faculty of Biology, University of Duisburg-Essen, Universitätsstraße 2, 45141 Essen, Germany

## Abstract

Despite decades of substantial research, cancer remains a ubiquitous scourge in the industrialized world. Effective treatments require a thorough understanding of macroscopic cancerous tumor growth out of individual cells. Clinical imaging methods, however, only detect late-stage macroscopic tumors, while many quantitative experiments focus on small clusters of cancerous cells in microscopic detail but struggle to grow full tumors *in-vitro*. Here, we introduce the critical scale-bridging link between both these scopes. We are able to simulate the growth of mm-sized tumors composed of 1.5 million *μ*m-resolved individual cells by employing highly parallelized code on a supercomputer. We observe the competition for resources and space, which can lead to hypoxic or necrotic tissue regions. Cellular mutations and tumor stem cells can lead to tissue heterogeneity and change tumor properties. We probe the effects of different chemotherapy and radiotherapy treatments and observe selective pressure. This improved theoretical understanding of cancer growth as emerging behavior from single-cells opens new avenues for various scientific fields, ranging from developing better early-stage cancer detection devices to testing treatment regimes *in-silico* for personalized medicine.

**Author summary:** Experimental and microscopy techniques are rapidly advancing biology and the observability of tissue. The theoretical understanding of tissue either focuses on a few cells or continuous tissue. Here we introduce the scale-bridging theoretical link that is able to model single cells as well as tissue consisting of millions of those cells, harvesting the power of modern supercomputers. We close the gap between single-cells and tissue through access to the full time-resolved trajectories of each cell and the emerging behavior of the tissue. We apply our framework on a generalized model for tumor growth. Tumor heterogeneity, as well as tumor stem cells are introduced, and the changes of behavior in response to cancer treatments is observed and validated.

## 1 Introduction

”The global cancer burden is estimated to have risen to 18.1 million new cases and 9.6 million deaths in 2018. One [..] in 8 men and one in 11 women die from the disease” [1]. Despite this ubiquity, cancer treatment is challenging as it is a highly specific disease whose progression severely depends on the affected organ, cause, and host. Tumor progression is based on many complex effects acting concurrently to facilitate the uncontrolled growth of some cells. Cell-internal processes, e.g. mutations up-regulating cell division or chemotherapeutic resistance, have a significant impact on the size, shape, and heterogeneity of the final tumor and strongly affect treatment response [2] and drug resistance [3]. Unfortunately, while such microscopic properties are of high importance, they are clinically poorly accessible. Hence treatment protocol and prognosis are inferred based on accessible macroscopic properties such as patient condition, tumor size as visible in MRI scans and biopsies. Bridging the scales between experimental single-cell findings and clinical data would significantly improve the understanding of cancer as an emerging property of its cellular composition [4]. This link would allow us to optimize chemo- or radiotherapeutic treatments. Ideally, personalized treatment strategies could be optimized by comparing the outcome of different treatment regimes.

One option for predicting tumor growth is leveraging the exponentially increasing computing capabilities of modern supercomputers. A crucial ingredient is the simulation parametrization, which is fueled by new microscopy techniques, and genomic tools that have made immense progress in the observation of cells, tissue, and the temporal evolution of those [5–7] as well as gene expression and mutations [8]. This already has driven modeling of tissue development and dynamics in the related fields of embryogenesis [9, 10], morphogenesis, tissue dynamics, homeostasis simulations [11–18], and tumor growth [19–24] simulations. These and other tissue modeling approaches paint an increasingly detailed picture, enabling predictive simulations that can be verified by experiments and *vice versa* for a large variety of biological phenomena [25, 26]. Still, the scope of prior cancer simulations is either the detailed description of individual cells or large numbers of cells as point-like agents or a coarse-grained description of tissue [27]. Nevertheless, both ends of the resolution range are necessary to map the complexity of tumor development, since both the individual cell as well as the macroscopic environment play a crucial role.

Here, we simulate the growth of a tumor inside a vascularized homogeneous tissue. Our cancer simulation considers competing single-cell effects leading to emerging tissue scale behavior. We introduce a computational microscope that enables access to time resolved trajectories of all included cellular properties, going beyond what is accessible in wet-lab experiments. We see that highly proliferative cells in a surrounding tissue form tumors of distinct shapes. The introduction of a nutrition dependent cell cycle leads to hypoxic and necrotic regions but also requires sub-cellular resolution to treat nutrient flow and other surface-based cell-to-cell interactions realistically. Tumor heterogeneity is incorporated by the mutation of cells into predefined cell-types reflecting driver mutations. Applying different treatment protocols of chemo- and radiotherapy with invariable doses leads to drastically different treatment outcomes and allows for a systematic scan of their effectiveness. Tumor stem cells (TSCs) [28] impact tumor heterogeneity development as well as treatment outcome in our model and can recover tumors after seemingly successful treatment. To handle the computational complexity of our model, we took advantage of modern supercomputer architectures and developed a new, highly parallelized software framework from scratch. Thus, we can model tissues up to a clinically relevant size of mm^3^ composed over a million individual geometry-resolved cells over time-scales up to a year at a temporal resolution of a minute.

## Model Description

Naturally, one has to balance model abstraction with its complexity, parameter availability, and computational costs. Here, we choose to focus to explicitly model cell geometry of both cancerous and regular tissue, mutations and cancer heterogeneity, nutrient availability from blood vessels (but no angiogenesis) and treatment and resistance development to both chemo- and radiotherapy in context of the host environment. More specific, our spatiotemporal multi-scale model describes the collective behavior of *O*(1 Mio) individual cells to simulate a macroscopic tissue of *O*(1 mm^3^)(= 1000^3^ voxel, cf. Figure 1 and SI:Movie1 and SI:Movie4). The model consists of three layers. The lowest layer is a *microscale 3D cellular Potts model* (CPM) layer which models the cells on a grid [14, 29–31]. The CPM is based on a Hamiltonian with local interactions. Modeling of cell-cell adhesion proportional to the cell interfacial surface, nutrient transport, and cell-to-cell signaling properties are explicitly dependent on the shape and surface of each cell, with each cell occupying around 10^3^ voxel corresponding to 1000 μm^3^. The CPM defines the mechanical properties of the cells, such as compressibility, volume constraints, and adhesive forces. The Hamiltonian energy function reads

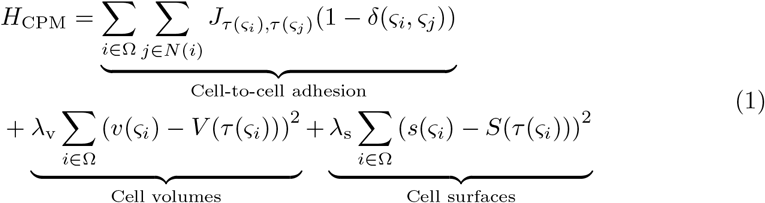

where Ω is the whole domain, and *N*(*i*) are the neighbors of voxel *i*. Further, *ς_i_* is the corresponding cell at voxel *i* of type *τ* and *ς_j_* is the corresponding cell at the neighboring voxel. The surface s of cells is calculated by a marching cubes algorithm, allowing an isotropic expansion in all directions. The energy function is implemented in a modular fashion allowing the addition of arbitrary additional energy terms e.g. for cell motility.

**Fig 1.**
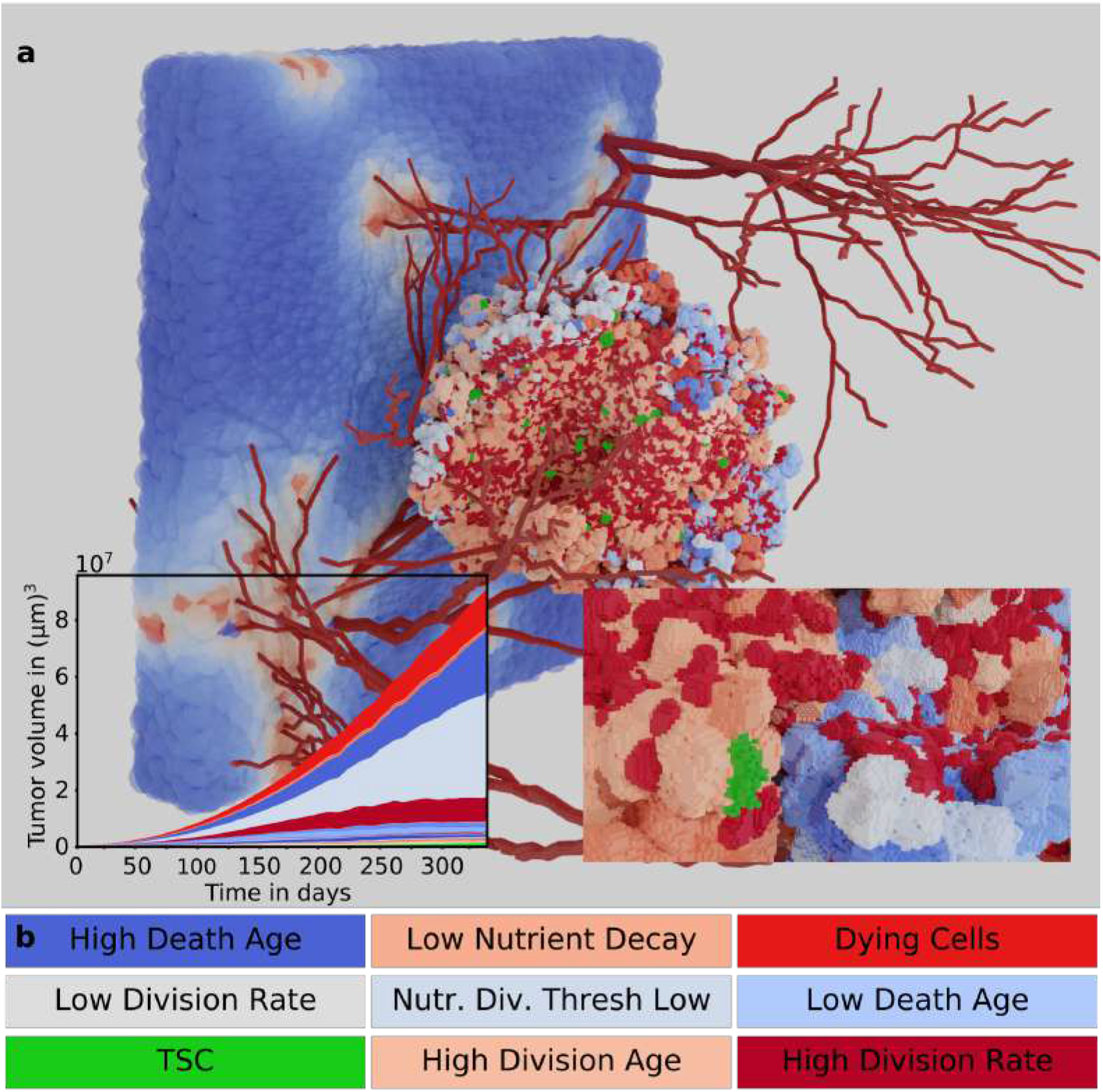
High-resolution Tumor Simulation: a) 1 mm^3^ tissue simulation with 1 μm resolution of single cells. Blood vessels (dark red) distribute nutrients (blue, white and red on sheet in background representing low, medium and high concentrations) that facilitate cell divisions and tumor expansion. Heterogeneous tumor growth (colors represent different cell types) results as an emergent behavior of nutrient dependant cell-division and -death as well as mutations. The inlays show growth of cell types over time (left), a zoom-in on the tumor surface highlights the *μ*m resolution. b) The color-coding of the majorly contributing cell types, each color indicates one cell type with its individual parameter-set, colors are used for the remainder of the figures (for all cell-types see SI:Figure 8).

Periodic boundary conditions reflect the behavior of an extended macroscopic tissue. The intermediate layer is a *mesoscale surface* layer, in which the diffusion of signaling compounds, nutrition, and chemotherapeutic drugs is realized by flux through the cell membrane to the adjoining neighbors of a cell. The transport is dependent on the local concentrations of each cell as well as a diffusion coefficient D. The flux 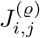 for a signal *ϱ* is defined by

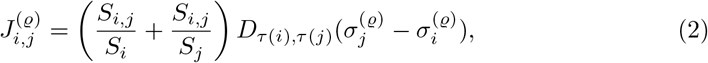

where *S_i_* is the surface from cell *i* and *σ_i_* is the signal value in cell *i* and *S_j_*, *σ_j_* from cell *j*, respectively. This is the arithmetic mean of the two surface fractions with respect to the common surface. The flux *J_i,j_* is subtracted from the signal of one cell and added to the other. Here, we distinguish between cells and fixed signal suppliers, such as blood vessels. For fixed signal supplier, the signal content is kept constant, i.e., the flux is neither subtracted nor added for those cells.

The top layer is a *macroscale agent-based* model that handles cell phenotype parametrization, cell internal signal processing, cell division, cell death, and mutations of cells. The cell type defines the parametrization of a cell, the subset of tumor cell types originates from the initial tumor type. Only a single parameter is changed for each tumor cell type. The surrounding tissue is initialized as a non-dividing and non-dying population. Cell division and cell death depend on the cell age, nutrient availability, cytostatic drug concentration, as well as division and death rates. Once cell death is induced, the cell is dying with its volume over time gradually lowered to zero. Mutations are possible events accompanying cell division, assigning a new phenotype to one of the daughter cells. Similarly, tumor stem cells (TCS) are implemented as cells with a slow cell cycle. Both mutations and TCS lead to tumor heterogeneity and affect treatment response and tumor progression and rejuvenation. Chemotherapy is implemented as a diffusive drug that suppresses cell division and is distributed via blood vessels. Radiotherapy introduces immediate cell death of a fraction of cells and globally reduces division rates proportionally to accumulated radiation exposure. The model parameters are largely based on experimental measurements (cf. SI:Table 2,3). It is possible to simulate arbitrary large simulation boxes with computational costs scaling with *O*(*L*^3^) with *L* measuring the box edge length. Most simulations use a cubic box with edge length of 320 *μ*m, with selected simulations using a box edge length of 1000 μm. A novel computational framework, *Cells In Silico*, handles the distribution of computing load resulting in super-linear speedup on large CPU-core numbers (*O*(10^5^)) on supercomputers.

For a detailed model, parameter, and framework description see SI:1 Methods. The published open source package of *Cells In Silico* in the NASTjA framework can be found at https://gitlab.com/nastja/nastja.

## Results

### Homogeneous Tumor Growth

Simulations of a small initial cluster of 35 cancerous non-mutating cells with high proliferative potential are carried out in a medium of surrounding tissue over a simulated time of one year at a one-minute time step (cf. SI:1 Methods, for simulation details). Over this time period, one can observe homogeneous tumor growth (i.e., composed of a single cell type) into the surrounding tissue. Nutrients represent a growth-limiting factor distributed by the blood vessels that could represent oxygen and or glucose. To better control the effects of nutrients, we distribute in a simulation box with an edge length of 320 μm blood vessels in a rectangular grid surrounding the initial tumor cells (cf. Figure 2 a)). In the simulations, the nutrients diffuse from blood vessels through the tissue, with each cell degrading the local nutrition concentration (cf. Figure 2 a) right). A gradient of nutrient concentration develops originating from the blood vessels (see SI:Movie2).

**Fig 2.**
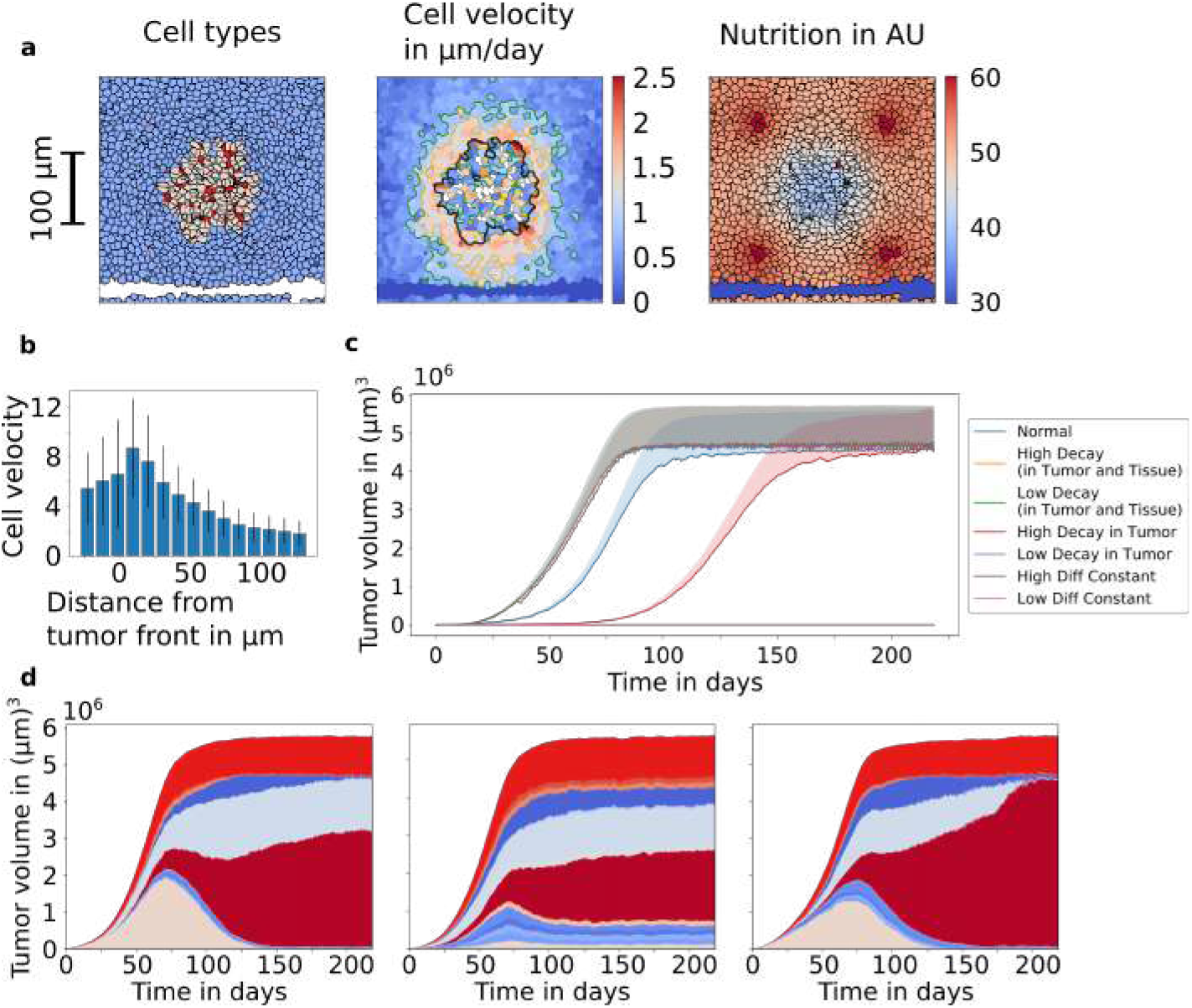
Buildup of the Model: a) 2D slices of tumor simulation with a rectangular blood vessel grid. Left to right: Coloring by cell types (Types see Figure 1), coloring by averaged cell velocities (black line indicates tumor outline), coloring by nutrient availability b) Cellular velocity in dependance of distance to tumor front, negative values are inside tumor. c) Dependence of the tumor growth rates on metabolism parameters. Parameter variations in the decay of nutrients and diffusion constant in the tissue and tumor change the growth rates of the tumors. The shaded area indicates the volume of dying cells. d) Tumor heterogeneity through mutation after cell division. A variation of the mutation rate results in different tumor compositions. The simulation on the left has medium, in the center a high mutation rate. On the right, the blood vessel configuration was changed for medium mutation rates. Colouring as in Figure 1.

As time progresses, the tumor invades the surrounding tissue. The tumor reaches a finite size once the simulated volume is entirely filled by cells. Growth becomes limited by compressibility, reaching a steady-state of proliferation and cell death. Tumor cells deplete nutrients at a higher rate leading to the formation of experimentally known intermediate states such as invasive fingers and hypoxic or even necrotic areas in the center of the tumor.

An up-regulated metabolism in tumor cells leads to a significantly decreased growth rate, while a down-regulated metabolism leads to faster growth of the tumor (cf. Figure 2 c)). Our analysis of cell velocities and cell displacement shows highly mobile or dynamic tumor cells at the boundary of the tumor. In contrast, the cell movements within the tumor and in the surrounding tissue, are much lower (cf. Figure 2 b)). Cell density and velocity have been associated with tumor invasion as well as jamming and unjamming transitions within a tumor [32].

### Heterogeneity

Primary tumors develop over long periods, and tumor internal heterogeneity arises from cells mutating during cell division. The limited inflow of nutrients leads to a competition of the cell phenotypes, and the partition into subpopulations indicates the fitness of the individual cell types.

We measure heterogeneity as:

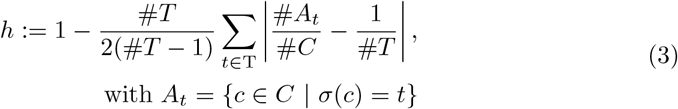

*T* are all cell types *C* are all cells *A_t_* are number of cells of type *t*, # denotes number. We run a set of simulations with different mutation rates and equal transition rates between the predefined tumor cell phenotypes. As visible in Figure 2 d), the final heterogeneity of a tumor strongly depends on the mutation rate. At low (every 200th division) and medium (every 20th division) mutation rates around day 70, the initial tumor cells dominate the tumor mass but get outcompeted with time as the total size of the tumor is stunted by lack of compressible surrounding tissue. At low mutation rates, cell types with a higher division rate and a delayed cell death, begin to dominate the tumor after day 70. Medium mutation rates lead to similar yet accelerated qualitative behavior. To observe the influence of the local environment, we increase the blood vessel density in the simulation leads to a more rapid preeminence of fast-dividing cells.

### Probing Treatment Regimes

Models to optimize chemotherapy dosage have been implemented and convincingly used as early as the 70s [33, 34]. Figure 3 a) depicts the response of simulated heterogeneous tumors to different conventional treatment schemes. We assume detection and onset of treatment of the tumor at day 110 until day 220. The drop in the total tumor size post-treatment until the final size at day 330 strongly depends on the treatment protocol. For different treatment protocols, the total dose of a therapeutic agent stays constant. It is redistributed into shorter peaks with higher concentrations and within the same time frame between days 110 to 220.

**Fig 3.**
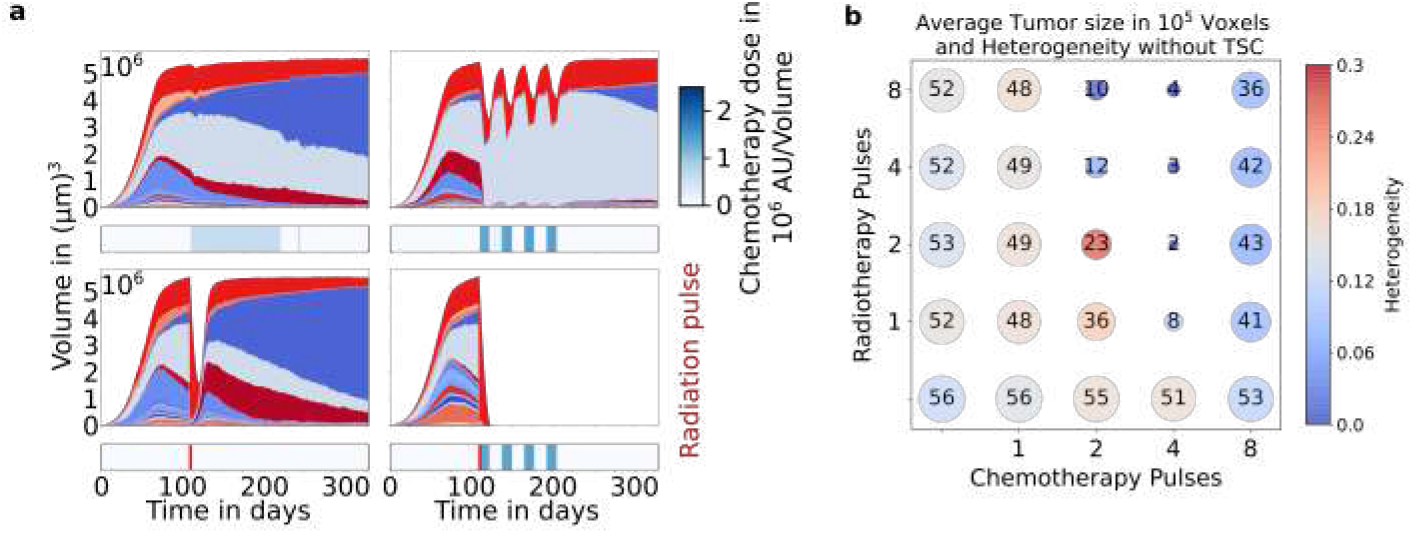
Treatment Response: a) Tissue size and composition response to different treatment regimes of constant accumulated doses of chemo- and radiotherapy. Colouring as in Figure 1. The treatment protocol of chemo- (blue) and radiotherapy (red) are depicted below the growth curve. b) Integrated tumor size (bubble size) post-treatment and tumor heterogeneity (bubble color).

Shorter pulses of chemotherapy show a greater effect than a uniform application, whereas a single strong radiotherapy pulse reduces the tumor size more drastically than multiple weaker pulses. Multiple pulses of radiation and chemotherapy shift the tumor composition towards a more homogeneous tumor by a cumulative adaptation through advantageous cell types surviving. This coincides with studies, which identify tumor heterogeneity as a driving force in treatment resistance [35]. Somewhat surprisingly, cell types with increased resistance to chemo- or radiotherapy are less favored than fast-dividing cells in the relapse post-treatment. Combinations of therapies using chemo and radiotherapy show a more significant effect on the tumor as one of the methods alone since the growth-inhibiting effect is two-fold. As clearly visible in Figure 3 b), treatment effectivity increases when going from one to four pulses of chemotherapy but then drastically decreases for eight pulses. For radiotherapy, the effect on the tumor also increases when dividing the dose into smaller pulses.

Thus, we can probe treatment regimes for a given tumor. We can systematically probe the treatment effects of different treatment regimes and combinations and judge effectivity based on tumor properties.

### Tumor Stem Cells

Tumor stem cells (TSC) are specialized cells within a tumor which through asymmetric cell divisions and a slower cell cycle, produce cancerous cells [28] and contribute to tumor rejuvenation as well as treatment resistance [36].

In our model, TSCs do not significantly change tumor size, growth rates, and final composition on unrestricted growth (see Figure 4 e)). The subpopulation of TSCs grows at a longer time scale due to a slower cell cycle (see Figure 4 e)) and are localized in small clusters but evenly distributed around the tumor (see Figure 4 d and SI:Movie3)).

**Fig 4.**
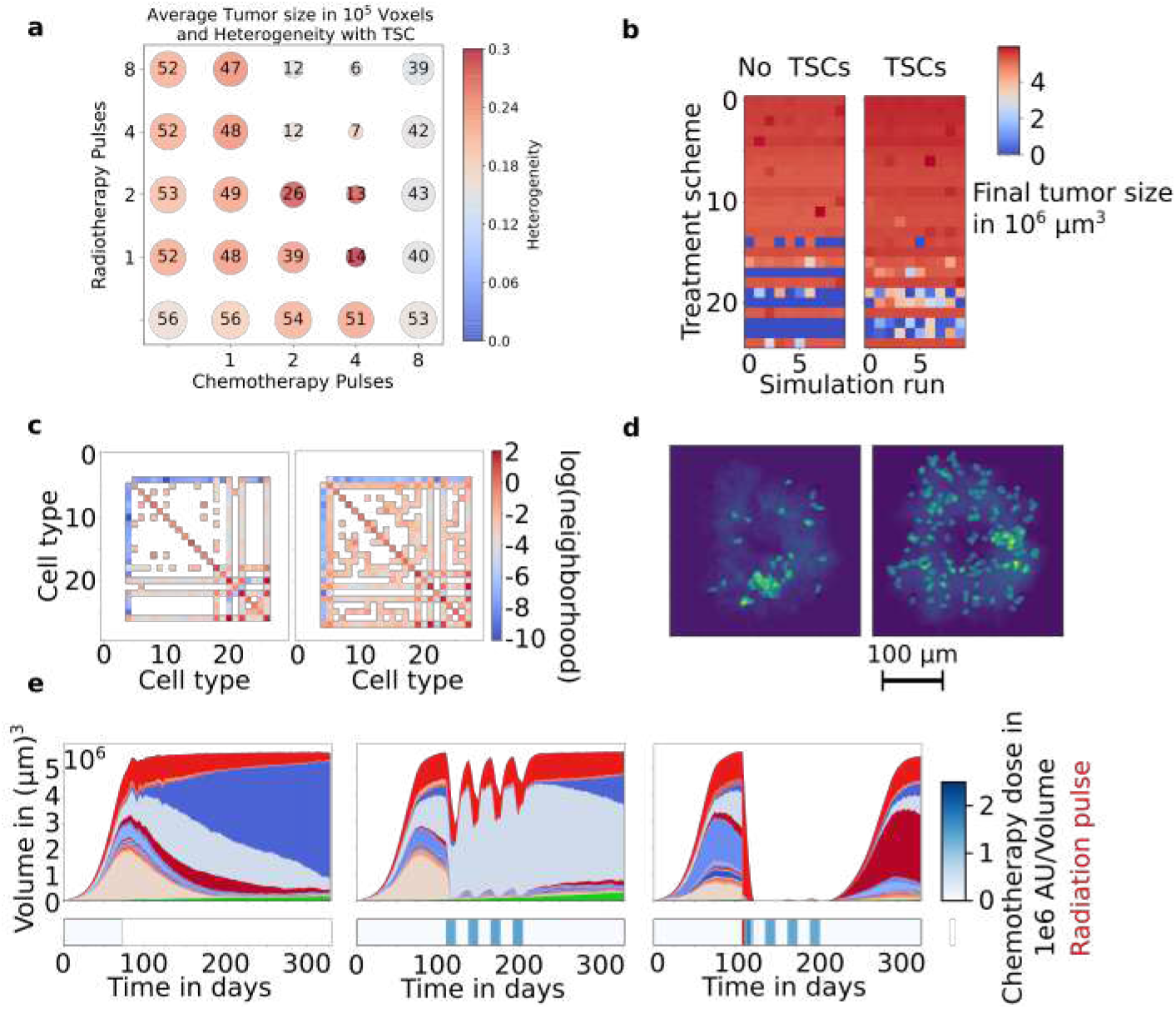
Tumor Stem Cells: a) Effect on final tumor size after treatment and tumor heterogeneity, errors as in Figure 3. b) The final tumor size of different treatment schemes (y-) and different simulation runs (x-axis) shows the stability of the simulation outcome. Colouring by tumor size at *t* = 325 days. c) Heatmap of the frequency of local neighborhood of cell types without (left) and with (right) TSCs. d) Localization of the tumor stem (in green) cells at *t* = 73 days (left) and *t* = 292 days (right), integrated over the entire volume. e) Response to different treatment schemes of chemo- and radiotherapy (Note regrowth bottom right). Colouring as in Figure 1.

The immediate treatment response is seemingly unchanged. Where subsequent chemotherapy in the heterogeneous tumor was able to suppress regrowth, TSCs can facilitate a subpopulation of cells to remain as a reservoir and trigger regrowth. TSCs lead to an increase in heterogeneity and boost the subpopulations of cell types that were suppressed. Figure 4 a) shows that TSCs can impact the final treatment outcome negatively by increasing tumor size as well as heterogeneity, especially in cases where treatment was successful in tumors without TSCs, see Figure 3 b).

In the heatmap in Figure 4 c), the type of a cell is compared to the types of its nearest neighbors. Diagonal elements are the major contributions, representing coherent clusters of cells of the same type. We find that TSCs lead to smaller cell clusters and a stronger mixing of cell types throughout the tumor, visible by increased off-diagonal elements. This is consistent with biologically observed behavior since tumors can regrow from a small number of remaining cells that are below the detection limit. [37]

### Stability

In order to provide predictions for the temporal development of tissue *in vitro* and *in silico* for clinical applications, knowledge about the statistics and robustness of the system development is essential. Variability in the simulation outcome is visible when running the same simulation with different random seeds. This reveals an impact of random and rare events on the macroscopic tumor development. The deviation in tumor size as well as heterogeneity in Figure 3 b) and 4 a) is neglectable, where the treatment weakly impacts the tumor size. Whereas for treatment schemes that drastically reduce the tumor size, the local surrounding and rare events have a more significant impact and lead to greater variability in the subsequent development. Figure 4 b) depicts the stochasticity of the final tumor volume after different treatment protocols and show increased variability in treatment schemes. In some cases, rare events can toggle between disappearance or relapse of the tumor post-treatment. TSCs lead to both less effective and more varying treatment outcomes.

## 2 Discussion

Our model highlights the possibilities of simulating emerging macroscopic tumor development resulting from microscopically explicitly shape-represented single cells. The high computing and data handling complexity can be mitigated via current-day supercomputing capabilities. The model makes it possible to test arbitrary ’what-if’ scenarios, unrestricted by experimental constraint, with direct control over all parameters of each individual cell. In our virtual tumors, we observe adhesion driven cell movements and nutrition-dependent heterogeneous tumor growth. We can show that different treatment plans strongly influence the final tumor cell type composition. We model cancer therapeutic agents in our system and show agreement with experimentally measured behaviors, reflecting growth curves [38] cf. SI:Table 3. Each simulation results in a fully spatio-temporally resolved trajectory, which allows tracing even single-cell events. Tumor stem cells introduce a source of treatment resistivity and are capable of facilitating a relapse of the tumor after seemingly successful treatment. Fundamental differences in the treatment response between tumors with and without TSCs were highlighted in this work, such as elevated intra-tumor heterogeneity and mixing. The stemness of a tumor has been experimentally associated with enhanced heterogeneity and treatment resistance [39, 40].

We observe that the tumor invasion is mostly driven close to the surface of the tumor and can investigate the changing tumor composition over time. In some simulation regimes, rare events influence not only details of the individual simulation but can influence the macroscopic, such as the resurgence of tumors post-treatment. This improved theoretical understanding of cancer growth as emerging behavior opens new research avenues. One could envision application in improved early-stage cancer detection by characterizing detectible early growth pathways. Once parameter sets for specific cancer types have been developed, such simulations could revolutionize clinical treatment via optimized, personalized medicine regimes *in-silico*.

## Supporting information

### 3 Methods

The model of *Cells In Silico* is composed of three different layers. Figure 5 depicts an overview of the layers and model parts that are acting on them.

**Fig 5.**
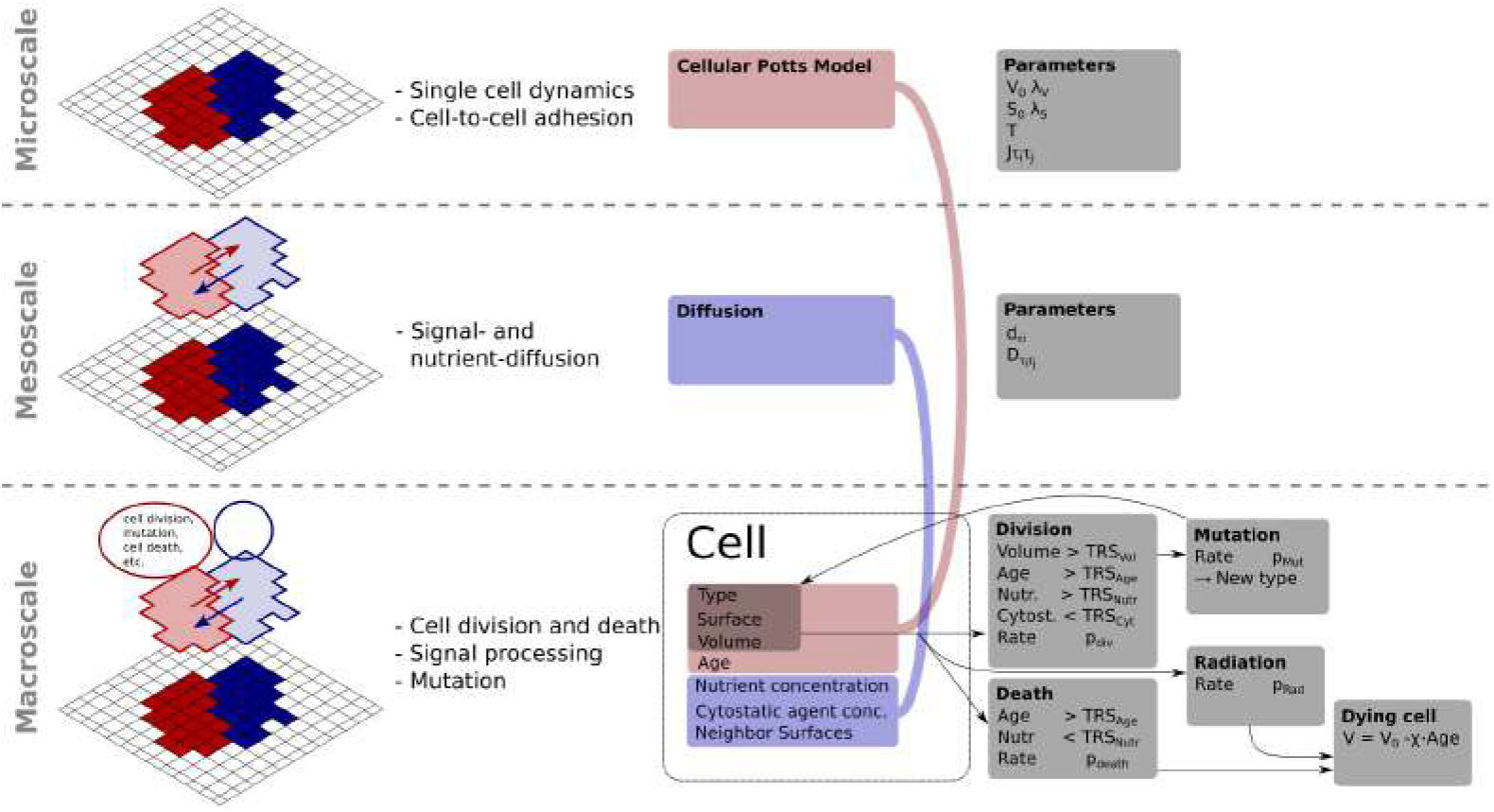
The schematic shows an overview of the simulation layers and actions of the agent-based model used in Cells in Silico.

Each layer represents a different length scale. Our model is based on a cellular Potts model acting on the microscopic layer, see Section 3.2. We add a general transport mechanism to our model. The diffusion of signals such as nutrients and drugs through the simulated area is modeled on the mesoscopic layer, which is described in Section 3.3. An agent-based model controls cellular events such as cell-divisions and cell deaths; this macroscopic layer is explained in Section 3.4. The parallel version of our model is implemented using the massively parallel NAStJA framework [41]. Besides, synchronization steps ensure a consistent state of the entire domain; these are the halo exchange as well as the local exchange of global cell properties is described in Ref. [42].

#### 3.1 Parallelization

The simulation area is discretized by a three-dimensional field containing a regular rectangular grid of voxels with a size of 1 *μ*m. Each voxel contains an integer value that denotes the cell ID. Voxels that have the same cell ID belong to an individual biological cell. The cellular Potts model (microscale) is acting on this data.

The whole domain is decomposed into small blocks, and these blocks are distributed to the different MPI (Message Parsing Interface) ranks, see Figure 6. The field in each block is enlarged by a halo layer, which overlaps with the fields of the neighboring blocks by one layer. This halo layer is updated for each block in every time-step to ensure consistency over the entire field. Besides the grid containing the cell IDs, each block holds global properties of all cells that are at least partially inside in the block, such as the cells volume *V*, surface *S*, see Figure 5.

**Fig 6.**
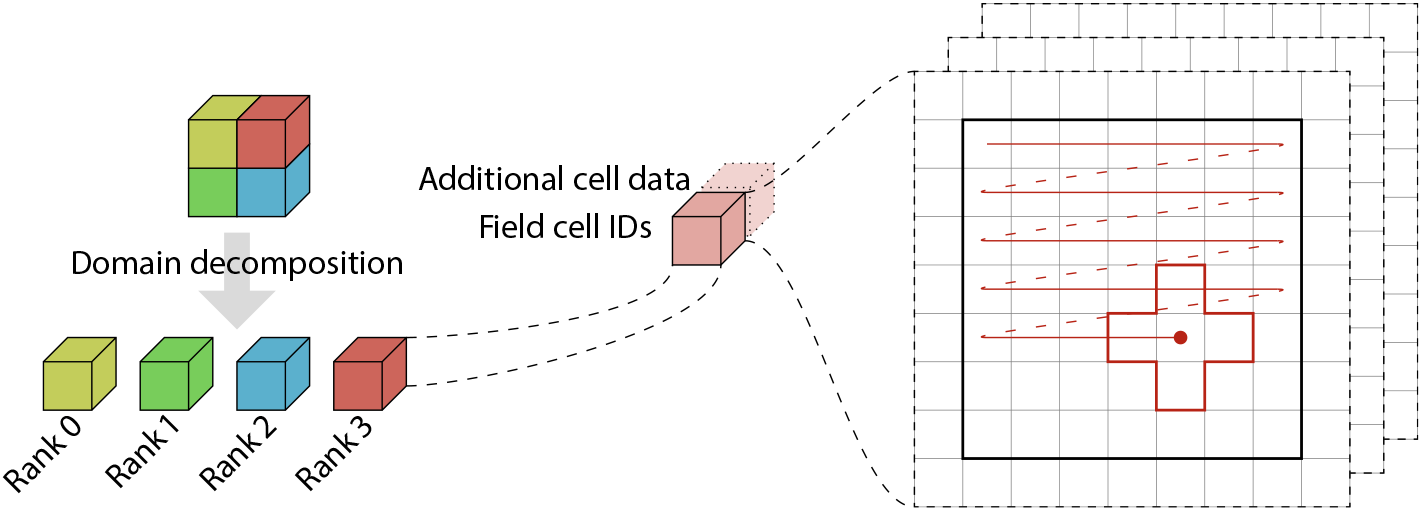
The domain is decomposed and distributed to MPI ranks. So each rank holds one block. Each block contains a field with the cell IDs and additional cell data. A field is a three-dimensional array on which the actions are performed.

In each time-step, a sequence of actions is executed independently on each MPI rank, see Figure 7. Actions that iterate over the field, such as the CPM propagation are called sweeps. Divisions and mutations are actions that are called every time step. After all sweeps and actions, the synchronization steps such as the halo and data exchange, as well as IO actions, are executed.

**Fig 7.**
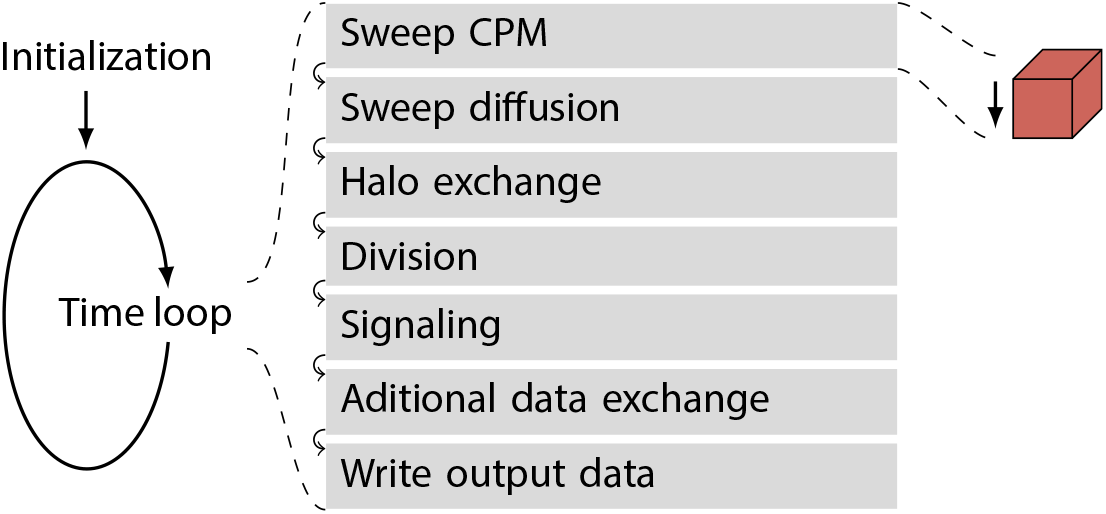
The actions in one time loop are structures in modules. Additional modules can be added depending on the simulated system.

**Fig 8.**
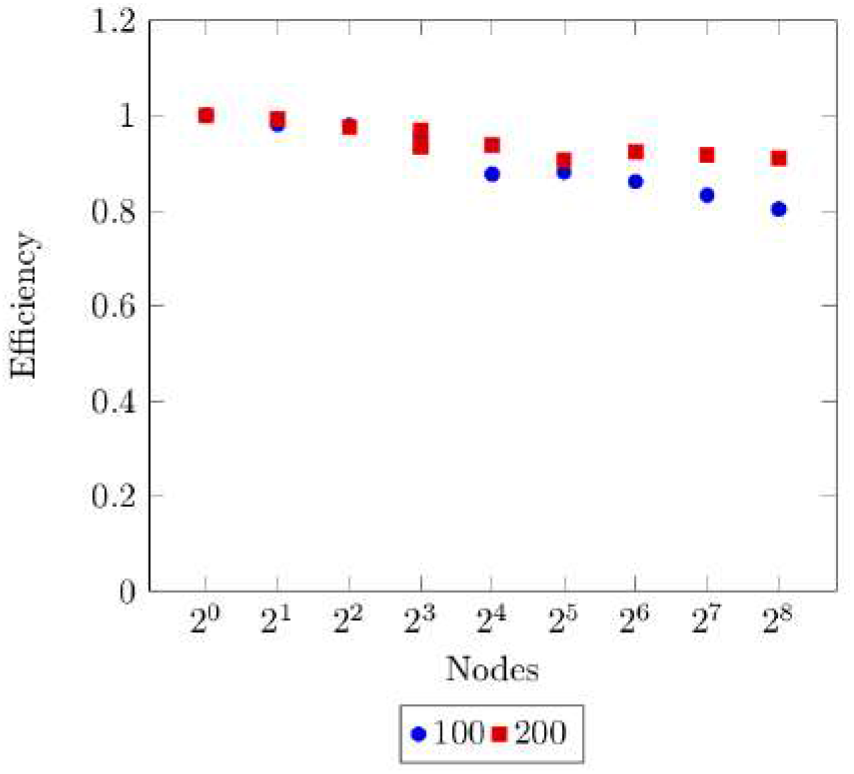
Weak scaling behavior. The efficiency of simulating large scale tissue on the Jureca supercomputer, using up to 256 nodes with 24 cores each. It is shown for a subdomain distribution with a 3D block edge length of 100 and 200.

Through a modular structure and parallelization of all sub-modules of the simulations we achieve excellent scaling behaviors up to a large number of cores 8.

#### 3.2 Microscale: Cellular Potts Model and Hamiltonian

The cellular Potts model was introduced by Glazier and Graner 1992 to simulate adhesion driven cell sorting [43].

It is based on a Potts model that describes integer spin states on a regular lattice, in both two and three dimensions. The temporal propagation of the system is performed by Monte Carlo sweeps over the field. Interactions are only possible between nearest neighbors and are accepted with the Metropolis criterion and the energy function.

The Hamiltonian is defined by the sum of several energy terms, reads,

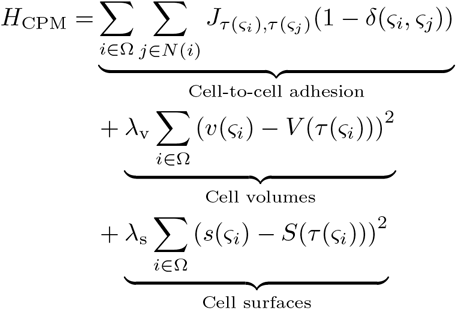

where Ω is the whole domain, and *N*(*i*) are the neighbors of voxel *i*. Further, *ς_i_* is the corresponding cell at voxel *i* and *ς_j_* is the corresponding cell at the neighboring voxel. Cell-to-cell adhesion is modeled by an energy contribution that is proportional to the shared surface (see 1.2.1)of different cells. *J* is the adhesion coefficient matrix giving the adhesion between two cells of types *τ*(*ς_i_*), *τ*(*ς_j_*), *δ* is the Kronecker delta, *v*(*ς_i_*) is the volume of cell *ς_i_*, *V*(*τ_i_*(*ς_i_*)) is the target volume of the cell type, λ_v_ is a coupling term regulating the strength of the volume constraint. *s*(*ς_i_*) is the surface of the cell *ς_i_*, *S*(*τ*(*ς_i_*)) is the target surface of the cell type, λ_s_ is a coupling term adjusting the strength of the surface constraint.

The system propagation in the cellular Potts model is based on random nearest neighbor interactions. The cell ID of a voxel can be changed to the cell ID of a randomly chosen nearest neighbor. Then, the energy difference Δ*E* of this local conformational change is calculated via the Hamiltonian energy function. Changes with negative energy differences are accepted, and positive energy differences have an exponentially decaying acceptance probability *p*_accept_, this is the Metropolis acceptance criterion.

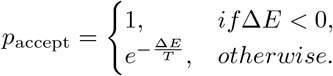

To perform the propagation of the entire grid, each voxel requires a uniform chance of sampling. For parallel execution, the system propagation has to be performed in every block, whereby it must be ensured that the changes in one block do not affect the changes in the neighboring blocks. In order to avoid the accumulation of errors at the block borders, a randomly chosen subset of 1/25 of all voxels that lie on cellular surfaces are sampled. The grid values are updated after such sweep over the whole block. Therefore all calculations are done on the grid state from the previous time step. Surfaces and Volumes of the cells are updated in the current block and communicated to all neighboring blocks. To ensure uniform sampling of the grid as well as avoiding runtime effects from earlier actions in a sweep, a visitor pattern is introduced that only allows changes in particular voxels, respecting the surface calculation metrics.

##### 3.2.1 Surface Calculation

The surface calculation metric is independent of the energy function. The calculation of the surface of objects on a cubic grid is not unique, depending on the chosen surface metric dependencies preferring some spacial directions that may occur, leading to anisotropies in the emerging structures. Traditionally, a Manhattan metric is used to calculate the surface in the cellular Potts model. The distance *d* between two points 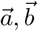 is defined by the sum of the absolute differences of their coordinates, 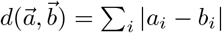. In two dimensions, this corresponds to counting the edges of the voxels and in the three-dimensional to counting the surfaces of the voxels. Under this metric, a unit circle has the same surface as a unit square. Likewise, in three dimensions, an ideal sphere of diameter *a* corresponds to a cube of edge length *a* after minimizing the surface. Particularly in the three-dimensional case, cell clusters tend to assume a cubic shape, when using the Manhattan-distance for the surface calculations, introducing a non-isotropic grid dependence in the model. In order to ensure a more isotropic sampling of the filed and to diminish grid artifacts, we use the marching cubes algorithm [44, 45]. The centers of eight voxels form the edges for the cube of the marching cube algorithm. Then we distinguish between all edges that have the cell ID that surface is calculated and all other cell IDs. Technically, we calculate the iso-surface for 0.5 by set the corners of the calculated cell ID to 1 and all others to 0. The surfaces of both algorithms are presented in Figure 9.

**Fig 9.**
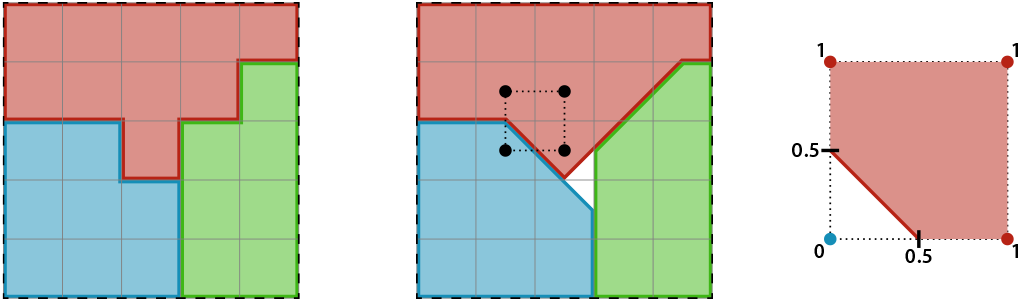
Manhattan surface calculation (left) and a two-dimensional representation of the marching cube surface calculation (middle). With a surface of 8 of the red cell(6.24 with marching cubes), blue and green cells have a surface of 6 (5.12 using marching cubes). The marching cubes are shifted at denoted by the black rectangle, i.e., for each voxel, there are four marching cubes in 2D and eight in 3D. On the right side is a detailed version of one marching cube, determined the surface for the red cell. The edges get the value 1 when it lies inside the red cell, 0 otherwise. The surface then is the 0.5 iso-line.

#### 3.3 Mesoscale: Signal and Nutrient Transport

The simulation considers the transmission and propagation of multiple substances, such as nutrients and drugs. We define a class of signaling, e.g., nutrient contents, of each cell 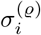, denoting the concentration of signal *ϱ* in cell *i*. Those represent either oxygen and glucose as nutrients for the cell, cell-to-cell signaling compounds or drugs. The diffusion of nutrients can be approximated by flow through the surfaces of the cells. Actions, such as cell-division, -death, and -mutations, may depend on these signals and nutrient contents.

##### Diffusion

The diffusion of signals between the cells occurs through the surface of these cells. We determine the shared surface *S_i,j_* for each pair of cells *i*, *j* with *i* ≠ *j*. The diffusion depends on the type of cells, so we define for each combination of types a diffusion constant *D*_*τ*(*i*), *τ*(*j*)_, *τ*(*i*) denoting the cell type of cell i. The flux 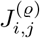 for a signal *ϱ* is defined by

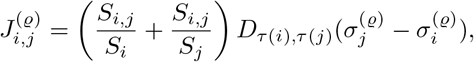

where *S_i_* is the surface from cell *i* and *σ_i_* is the signal value in cell *i* and *S_j_*, *σ_j_* from cell *j*, respectively. The first bracket is the arithmetic mean of the two surface fractions with respect to the common surface. The flux *J_i,j_* is subtracted from the signal of one cell and added to the other. Here, we distinguish between cells and fixed signal suppliers, such as blood vessels. For fixed signal supplies, the signal content is kept constant, i.e., the flux is neither subtracted nor added for those cells.

##### Decay

Metabolic processes take place inside the cells. We used a simple mesoscale model in which the signals are changed relative to their value,

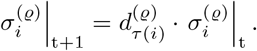

Where 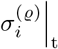 is the signal *ϱ* in cell *i* at time *t* and 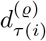 is the relative change of the signal *ϱ* depending on the type of cell *i*.

#### 3.4 Macroscale: Agent-based Method

On the macroscale, cell attributes such as the cell age, the signal level, cell type, etc. are used to generate actions based on these values. The parameters can be linked from cell biological experiments and simulations (cf. Table 2).

**Table 1.**
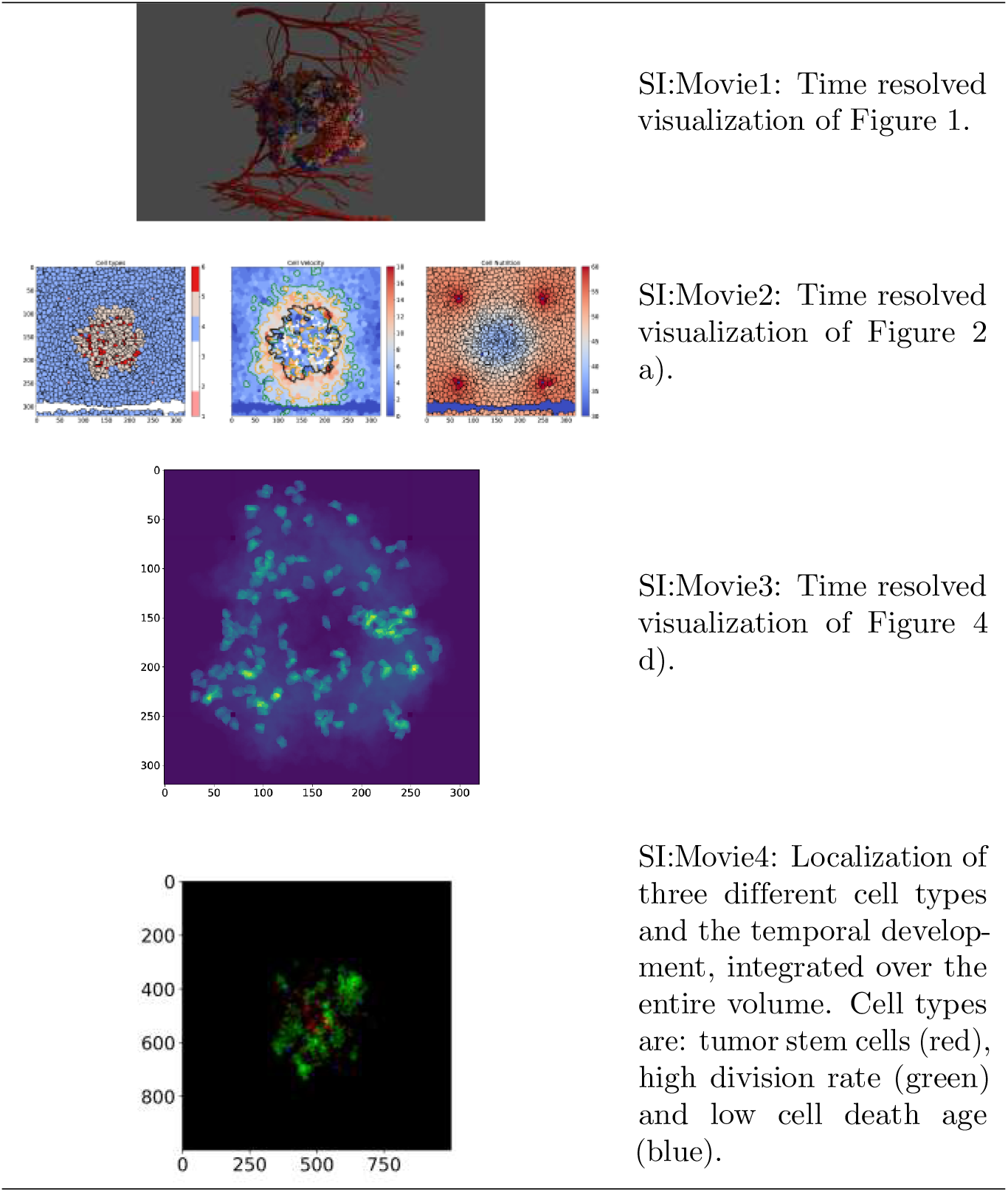
Supplementary Movies

**Table 2.**
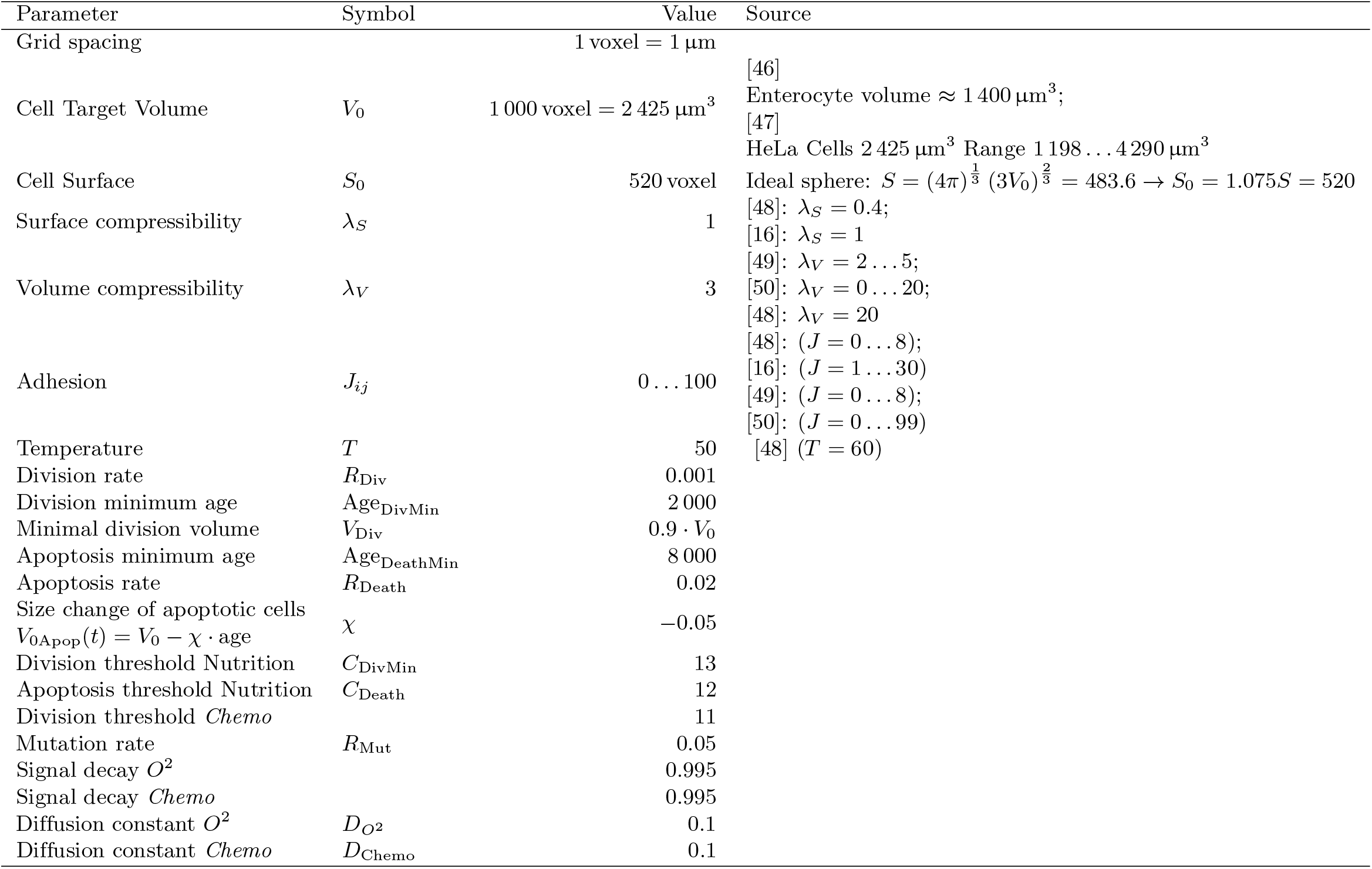
Model parameters that were used in the simulations.

**Table 3.**
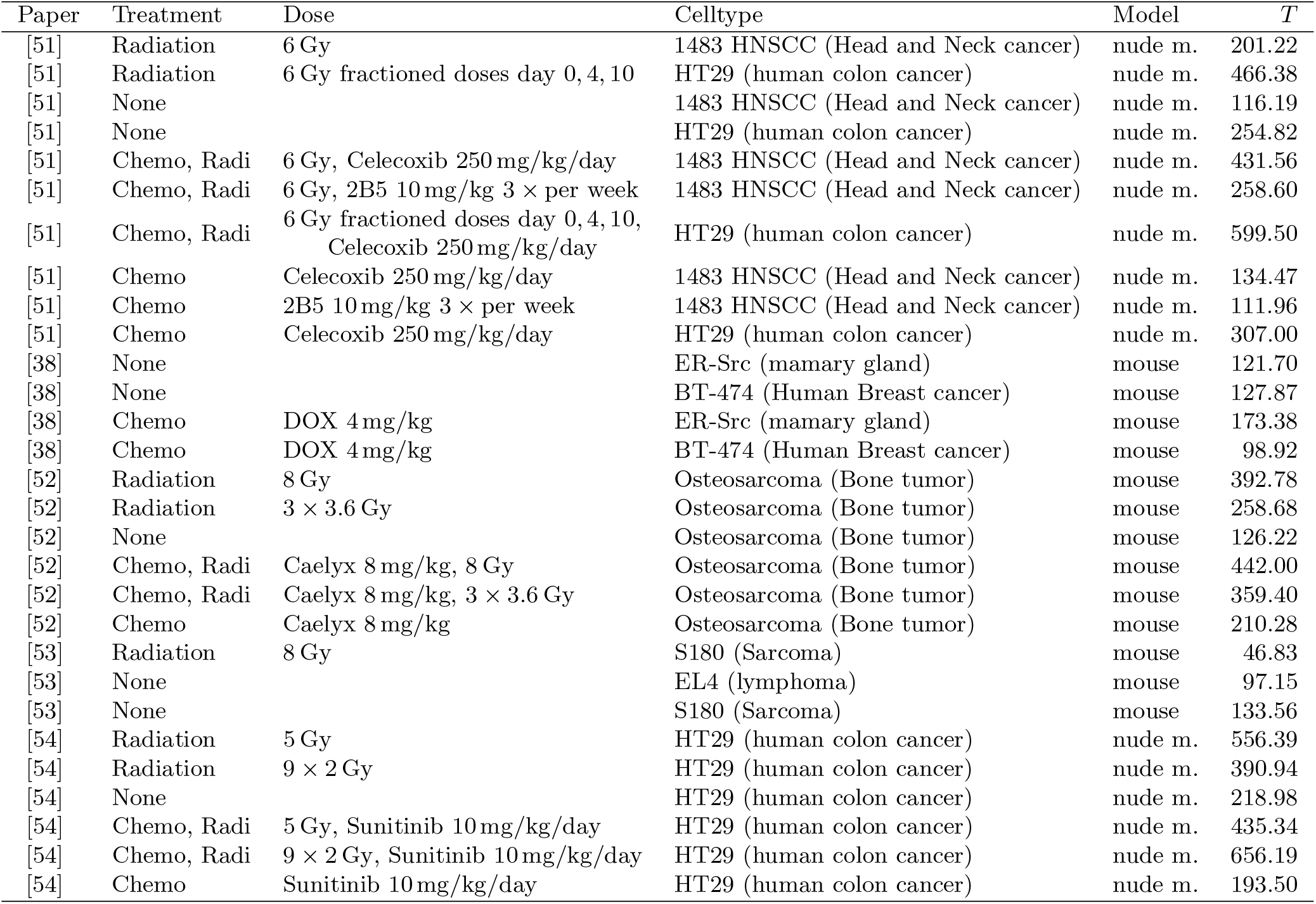
Fitted tumor growth time constants from various *in vivo* experiments, mostly on mouse models. Time constants *T* (in hours) were fitted for tumors without and with treatment (chemotherapy and/or radiotherapy). Through manual literature mining, experimental parameters of growth rates of tumors were extracted. Exponential growth was fitted to control tumor growth and to treated tumor growth.

##### Cell types

A cell type is assigned to each cell, which determines the parametrization and phenotype of that cell. The cell type defines the characteristics of the individual cells, i.e., the target volume *V*_0_, the target surface *S*_0_, and the thresholds TRS_Vol_, TRS_Age_. In that way, not each cell has to be individually parameterized. Cells that divide usually generate two new cells of the previous cell type. Cell types allow classification of each cell in the simulation as well as the tracking of cell type sub-populations. Through the definition of a predefined set of cell types and mutation between those types, the parameter space is controlled, and the accumulation of purely favorable traits in a single cell type is prevented. Each cell type only has a single variation in respect to the initial tumor cells. It is possible to define an arbitrary number of cell types for different use cases. Here we define a set of 27 cell types for the heterogeneous tumor growth (see Figure 13)

##### Blood vessels and solid

We introduced a subset of cells that is not participating in the spacio-temporal propagation via the cellular Potts model. Those cells are solid structures, which can model blood vessels or the extracellular matrix. They are able to participate in cell-to-cell signaling and may act as sources for signals.

##### 3.4.1 Actions of the Agent-based Model

###### Cell division

In each time-step, each cell is checked for cell-division. Whether a cell divides depends on several internal and external factors. Division conditions are:

- Volume above a threshold *V* > *V*_Div_ = 0.9 · *V*_0_
- Nutrition above a threshold *C*_DivMin_
- Age above a certain threshold Age_DivMin_
- Comparing random number ∈ [0, 1] with Division Rate *R*_Div_
- Chemotherapy content below a threshold TRS_Ch_

If all conditions are met, a random plane through the cell centre is chosen. The cell is split along that plane. Due to the parallelization of the model, the cell-division has to be communicated to neighboring blocks. The cell is split, and the two arising cells are reinitialized, measuring surface and volume. One keeps the cell ID of the mother cell while the other receives a new cell ID. After a cell-division, the cellular age is set to zero. Post division both cells expand due to pressure by the volume and the surface energy term. Specific cell types can also be excluded from cell-division, such as the surrounding tissue in our simulations.

###### Cell death

Cell death conditions are:

- Nutrition below a threshold *C*_Death_
- Age above a certain threshold Age_DeathMin_
- Comparing random number ∈ [0, 1] with death Rate *R*_Death_
- If all above conditions are not met comparing random ∈ [0, 1] with default death Rate *R*_Death_/1000 to account for natural cell death

Cell death is induced by changing the cell type of the cell to a dedicated cell type that describes dying cells. For this cell type the goal volume in the Hamiltonian is changed over time *V*_0Apop_(*t*) = *V*_0_ – *χ* · age, effectively lowering the volume of the cell to zero voxels. Once the cell reaches *V* = 0, the cell is deleted.

###### Mutation

After cell-division, the two daughter cells are reinitialized. If a mutation event occurs (mutation Rate *R*_Mut_), one of the daughter cells is initialized with a randomly chosen cell type. The range of cell types that can be chosen is predefined. The transition matrix between all cell types can be defined so that the transition probabilities between cell types varies. Here, we use a constant transition probability.

### 4 Simulations

#### 4.1 Time-step

The cellular Potts model is a discrete-time Markov chain. Incremental nearest neighbor interaction Monte Carlo steps model the fluctuations and movements of cell membranes.

We observe initially exponential growth of the *in-silico* tumor when modeling the free growth of a tumor in an environment with sufficient nutrients and without treatment. We find that a size doubling time of *T* = 8552 MonteCarloSweeps(MCS) in our model corresponds to an untreated tumor with an experimentally determined doubling time of 150 h (see Figure 10). Comparison with growth observed *in-vivo* (cf. Table 2) we assume exponential growth of the tumor volume *V*_Tumor_(*t*) = exp(ln(2) · *t/T*) + *c* and derive *T* = 150 h from the experimental data for a non-treated tumor. Therefore 1 MCS equals 1.05 min and 1 kMCS = 0.73 days.

**Fig 10.**
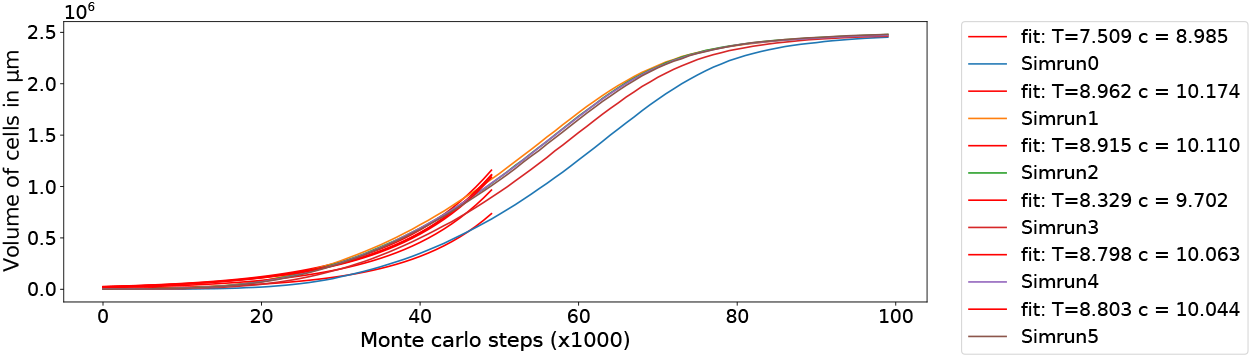
Growth of a tumor in surrounding tissue without external influences is used to define the timestep. Multiple simulation runs with different seeds are shown, and the time constant of the growth is fitted.

**Fig 11.**
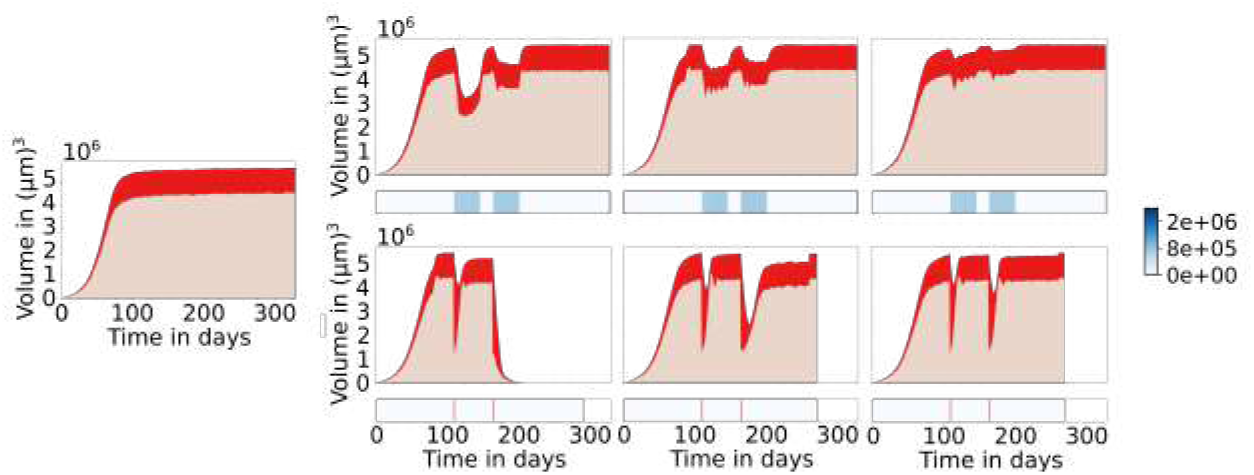
Tumor size response of a single cell type tumor of varying resistivities. From left to right: No Treatment, low, medium and high resistance against treatment. Chemotherapy schemes on the top row and radiation therapy schemes on the bottom row.

**Fig 12.**
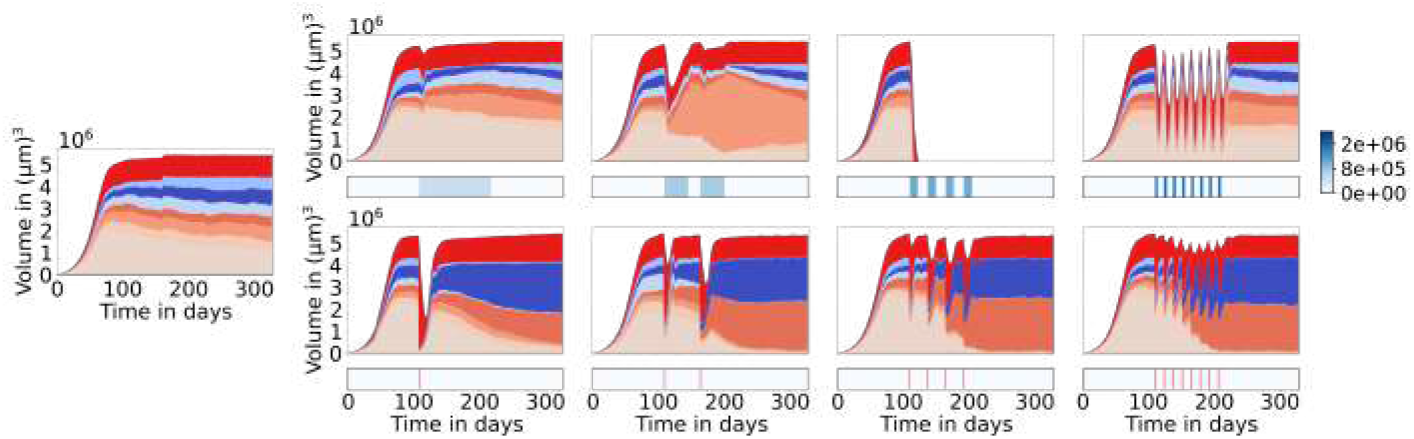
Treatment response to a heterogeneous tumor. The cell types are a different subset of cells than used for heterogeneity simulations in the main text, cell types only differ in treatment resistivity. Chemotherapy schemes on the top row and radiation therapy schemes on the bottom row.

**Fig 13.**
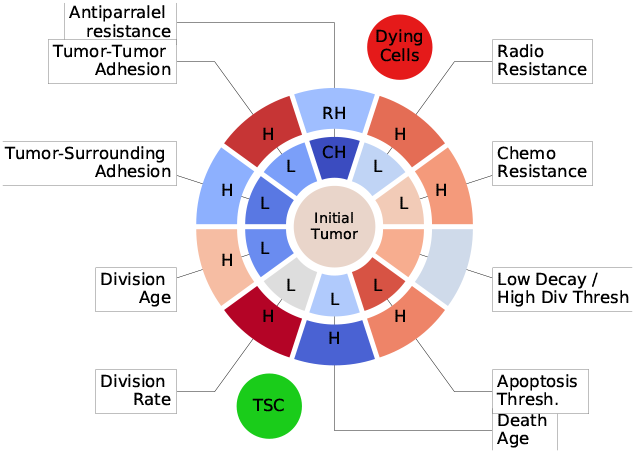
Colour coding of all cell types. The inner circle holds cell types with down-regulated parameters, while the outer circle holds up-regulated parameters.

**Fig 14.**
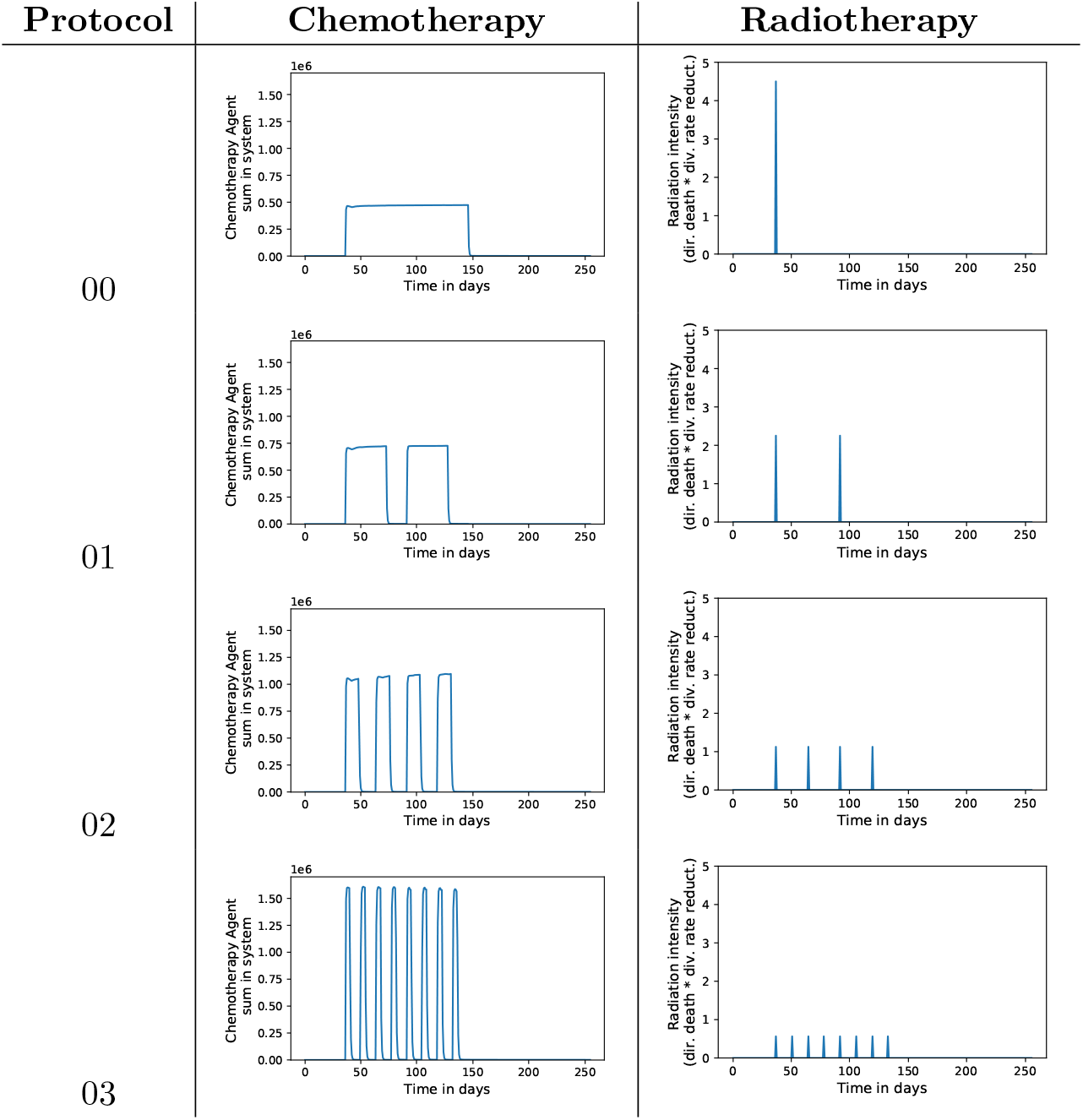
Overview over the treatment schemes of chemo- and radiotherapy.

#### 4.2 Simulations

General simulation parameter can be found in Table 2.

##### 4.2.1 Initial Simulations

###### Treatment of one cell type

Homogeneous tumor growth in a non-dividing surrounding tissue. Simulations are conducted in a box with an edge size of 320 μm. Variation of treatment resistivity and resulting tumor response of a two-pulse treatment with radio- and chemotherapy, respectively.

###### Treatment of resistivity heterogeneous cell type

Heterogeneous tumor growth in a surrounding tissue with a box edge size of 320 μm. Variation of treatment resistivity and resulting tumor response of a two-pulse treatment with radio- and chemotherapy, respectively.

##### 4.2.2 Heterogeneity

In tumor development, cells acquire more mutations until one mutation provides properties that enable the tumor to expand and spread. This mutation process continues during the whole lifetime of the tumor. Hence, a tumor does not consist of just one cancerous cell type, but a variety of cellular phenotypes that compete over resources and complicate tumor treatment since the effect of therapies differs between the types. Therefore tumor heterogeneity is a crucial factor in planning cancer treatments [35].

We implemented the mutation of cells into our model by introducing the possibility of a change of cell-type of one of the daughter cells after division. These mutations are reflecting one ore multiple somatic mutations that affect the behavior of the cell and lead to an altered behavior. Each of the predefined cell types represents one cellular phenotype and has exactly one parameter up- or down-regulated. The transition rates are constant from and to every cell type. Cell to cell adhesion, cell-division ages and nutrient thresholds, cell death age and thresholds as well as nutrient uptake, and division and death rates were altered.

##### 4.2.3 Treatment

Since cancer is such a versatile condition, many different forms of treatment exist with chemotherapy, radiation therapy, and immunotherapy being the most prominent. In a growing tumor, the balance of cell-division and cell death is shifted towards cell-division, leading to an uncontrolled expansion of the tumor. The goal of classical cancer therapies is to shift that balance towards the favor of the cell death rate, leading to depletion, and finally, the vanishing of cancerous cells. Chemotherapy and radiotherapy aim at damaging cells and therefore altering the division and apoptosis rates for all cells. Since cancerous cells have a significantly shorter life-cycle and weakened repair mechanisms in comparison to healthy tissue, the damage affects tumor cells to a greater extent than the surrounding healthy tissue. Therefore a specific dosage and protocol have to be used to do enough damage to shrink the tumor while keeping the damage to all other parts of the organism to a minimum. There are different chemotherapy protocols for each type of cancer, most opting for a pulsed administration over a longer time in order to maximize efficiency and reduce the development of resistance. In our model, *chemotherapy* is realized by a cytostatic agent that is delivered from the blood vessels and diffuses into the tissue at defined times. Depending on the local drug concentration cells down-regulate their division.

*Radiation therapy* is realized by a down-regulation of the cell-division rates and cell death initiation with a set probability for each tumor cell. The effect of radiation is homogeneous over the whole simulated area, since the radiated areas are clinically much bigger than the simulated domain. Four different treatment plans are compared for each, radio- and chemotherapy and applied alone, and in combination, to the heterogeneous tumor simulation.

##### 4.2.4 Up-scaling

An interesting question is how the tumor development and treatment response depend on the tumor age and size. We take advantage of the scalability of our simulation framework and scale up the simulations by a factor of 27 so that the simulation covers 1 mm^3^ of tissue. The surrounding tissue is initialized to be densely vascularized.

In the tumor growth we see due to the bigger size of the tumors, more complex structures arising. The tumor grows into the direction of the closest blood vessels and even divides up into smaller compartments. Holes in the tumor emerge, and apoptotic regions are repopulated with the surrounding tissue.

## Acknowledgments

We thank Jens Elgeti as well as Ines Reinartz for the support and comments that greatly improved the manuscript. The authors gratefully acknowledge the Gauss Centre for Supercomputing e.V. (www.gauss-center.eu) for funding this project by providing computing time through the John von Neumann Institute for Computing (NIC) on the GCS Supercomputer JUWELS at Jülich Supercomputing Centre (JSC). This research was supported by the Helmholtz Impuls- und Vernetzungsfonds. This work was partly performed on the supercomputer ForHLR funded by the Ministry of Science, Research and the Arts Baden-Württemberg and by the Federal Ministry of Education and Research.

